# Brain energy metabolism is optimized to minimize the cost of enzyme synthesis and transport

**DOI:** 10.1101/2022.11.14.516523

**Authors:** Johan Gustafsson, Jonathan L. Robinson, Henrik Zetterberg, Jens Nielsen

## Abstract

Energy metabolism of the brain is poorly understood partly due to the complex morphology of neurons. Here we used metabolic models that estimate costs of enzyme usage per pathway, enzyme utilization over time, and enzyme transport to evaluate a paradigm that suggests that brain energy metabolism is optimized to minimize enzyme synthesis and transportation costs. Our models recapitulate known metabolic behaviors and provide explanation for the astrocyte-neuron lactate shuttle theory.

## Main text

To sustain the energy demand of the brain, its cells combine glycolysis and mitochondrial respiration in a complex pattern that varies across cell types, brain regions, and degrees of activity. Specifically, it has been observed that 1) regions of the brain consume less oxygen per glucose molecule at high activity^1–4^; 2) the use of lactate as substrate to fuel the TCA cycle is important to sustain brain function in health and disease^5,6^; and 3) astrocytes tend to produce lactate that is consumed by neurons, a phenomenon known as the astrocyte-neuron-lactate-shuttle (ANLS) theory^4,7^. However, the underlying drivers behind these behaviors are largely unknown. Here, we present a theory that the brain has optimized its metabolism to minimize energy expenditure on maintaining functional ATP production pathways, because of frequent exposures to starvation during evolution.

The physiology of neurons differs substantially from that of most other cell types in that they have long extensions known as dendrites and an axon, where the latter can reach a length of up to 1 meter^8^ (Fig. 1A). Since most proteins are generated in the soma they need to be transported to their final locations and back for degradation^9^, which requires additional ATP costs to maintain the proteome. A large portion of the ATP usage in neurons occurs at the synapses^10^, where it is generated locally, creating a high demand at these sites for enzymes involved in ATP production pathways.

**Fig. 1:**
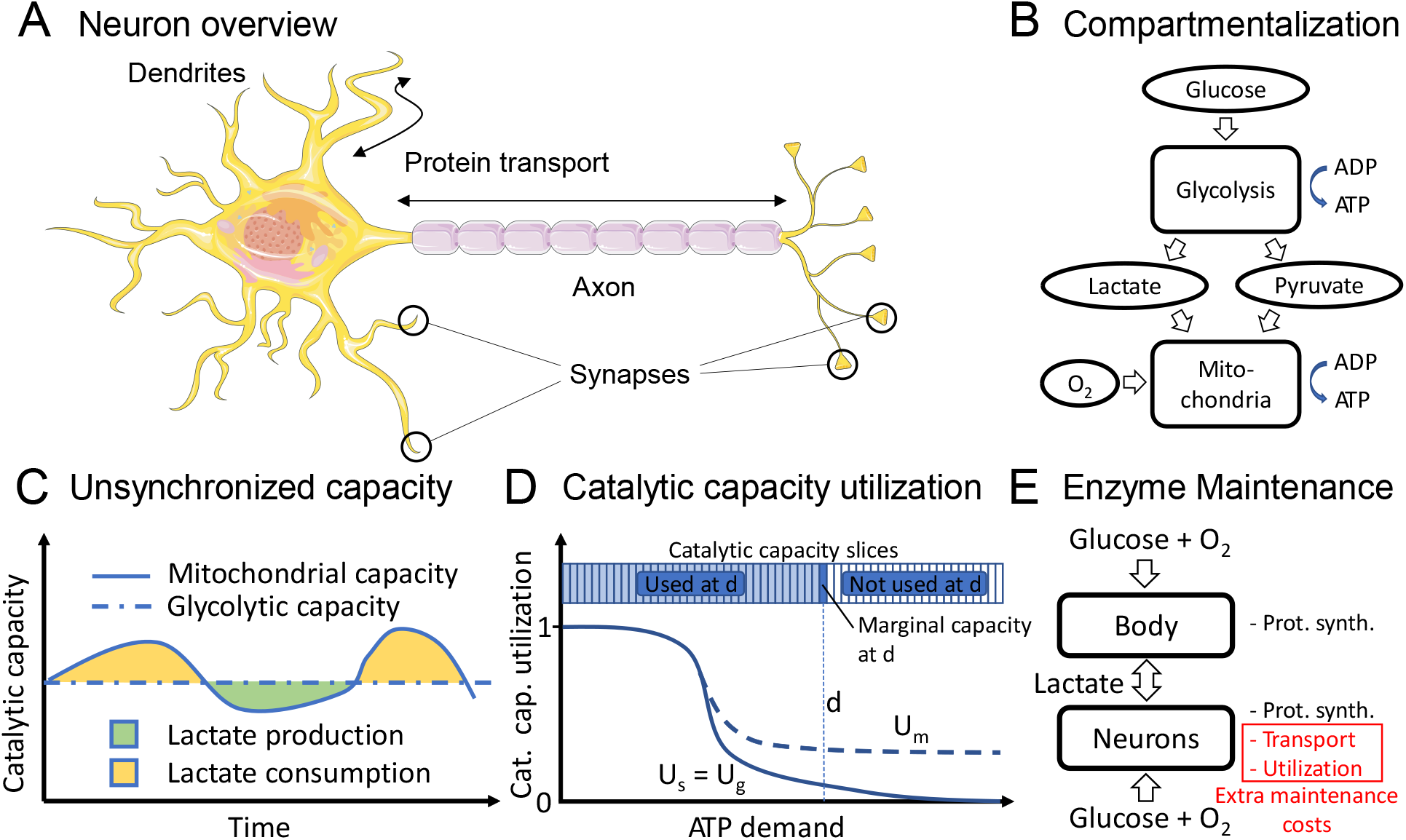
Neuron energy metabolism. A. Neuron overview. Enzymes must be transported to distant compartments of both the axon and dendrites, which requires additional ATP to maintain fully functional metabolic pathways there. B. Energy metabolism compartmentalization of glycolysis and mitochondrial respiration. Glycolysis can operate independently of mitochondria by production and export of lactate. Mitochondria can also in turn operate independently of glycolysis, if lactate is available. C. The availabilities of glycolytic and mitochondrial enzymes are not synchronized over time. The mobility of mitochondria leads to an uneven presence of mitochondria over time, causing the availability of enzymes catalyzing the TCA cycle and OXPHOS to vary over time at a certain position. Lactate availability enables additional ATP generation during time periods when the abundance of mitochondrial enzymes is high compared with that of glycolytic enzymes. D. Catalytic capacity utilization. The energy demand at a certain physical location in the neuron, for example at a synapse, varies over time. To manage the high energy demand during peaks, the enzyme availability must be high, although the additional catalytic capacity supplied to address these peaks will only be utilized at a fraction of the time. The catalytic capacity needed to produce ATP is here divided into infinitely small slices, where each slice represents the marginal capacity needed to sustain a certain ATP demand (d in the figure). The marginal catalytic capacity required to sustain a certain ATP demand d will only be used at time points when the ATP demand >= d and thus only a fraction of the time. This fraction of time is defined as the catalytic capacity utilization, and the cells need to always be able to have catalytic capacity to meet these peak demands. When the ATP demand is increasing, the catalytic capacity utilization will therefore decrease and lead to large maintenance costs per ATP produced. U_s_, U_g_, and U_m_ represent the catalytic capacity utilization for static, glycolytic, and mitochondrial enzymes, respectively (see main text). E. Enzyme maintenance ATP costs from a whole-body perspective. The ATP costs for maintaining the enzymatic capacity for ATP synthesis are in most cells mainly associated with enzyme synthesis and degradation but transportation and low utilization add extra maintenance costs in neurons per produced ATP. To optimize the use of glycolysis and mitochondrial respiration between the neurons and the rest of the body, lactate can be transported between them via the blood, however at a limited rate.

The main ATP generation machinery can be grouped into two processes: glycolysis, which is performed by cytosolic enzymes, and mitochondrial respiration, which is performed in the mitochondria and includes the TCA cycle and oxidative phosphorylation^11^ (Fig. 1B). Glycolysis can operate independently of mitochondrial respiration by converting the end-product pyruvate to lactate followed by export; likewise, mitochondrial respiration can function independently from glycolysis if provided a fuel such as lactate or ketone bodies.

Mitochondria are known to be mobile and move towards locations with high ATP demand^10^, which leads to varying mitochondrial capacity over time at a given location (Fig. 1C). Although glycolytic enzymes are capable of moving in response to hypoxia^12^, we consider this effect to be limited in comparison to mitochondrial mobility and therefore assume the glycolytic capacity to be stationary and constant over time. At a given intracellular position, there will therefore be time periods with relatively lower mitochondrial capacity, in which lactate can be exported to maximize usage of glycolytic enzymes, and likewise periods with higher mitochondrial capacity, which can only be fully utilized if lactate or a similar fuel is available.

The ATP demand at a specific physical location in the neuron can vary substantially over time, especially at synapses^10^, and to meet peak ATP demands, a matching ATP production capacity must be available. For stationary enzymes, the high capacity must always be present, even though the full capacity will only be used a small fraction of the time. We define the *catalytic capacity utilization U(d)* (henceforth just utilization) as the fraction of the time the catalytic capacity required to support the ATP demand *d* is actually needed, where the *static utilization U_s_(d)* is the utilization of stationary enzymes (Fig. 1D, Note S1). We further define *U_g_(d)* as the utilization of glycolytic enzymes, which due to low mobility is assumed to be equal to the static enzyme utilization, and *U_m_(d)* as the utilization of mitochondrial enzymes. Since mitochondria are known to be mobile, the utilization of mitochondrial enzymes is likely higher than the static enzyme utilization since some of the mitochondrial enzymatic capacity can be used at other physical locations during non-peak time periods. This makes it possible to share enzyme maintenance costs across locations.

We envision that the catalytic capacity allocated to produce ATP can be divided into infinitely thin slices, where each slice can produce an equal amount of ATP and represents the marginal catalytic capacity required to sustain a certain ATP demand (Fig. 1D). Each slice can then be allocated to either glycolysis, mitochondrial respiration, or both, depending on what combination generates the least enzyme maintenance cost from a whole-body perspective (Fig. 1E). The enzyme maintenance energy per synthesized ATP is higher at synapses due to extra costs from enzyme transportation and low utilization. We define the extra maintenance cost as the “ Extra ATP Maintenance Cost per ATP produced” (EAMCA), where the unit of EAMCA represents the enzyme synthesis and degradation cost of glycolysis per produced ATP (without taking transportation or utilization into account).

Genome-scale metabolic models describe all the metabolic reactions of a cell^13,14^. We constructed such a model incorporating enzyme usage information, which has proven useful for explaining metabolic behaviors in for example muscle cells^15^ and tumors^16^, and used that information to estimate ATP maintenance costs for enzyme transportation and utilization (Fig. 2A). Enzyme half-lives also affect the maintenance costs of enzymes but were found similar across the main pathways and were therefore not included in our model (Fig. S1).

**Fig. 2:**
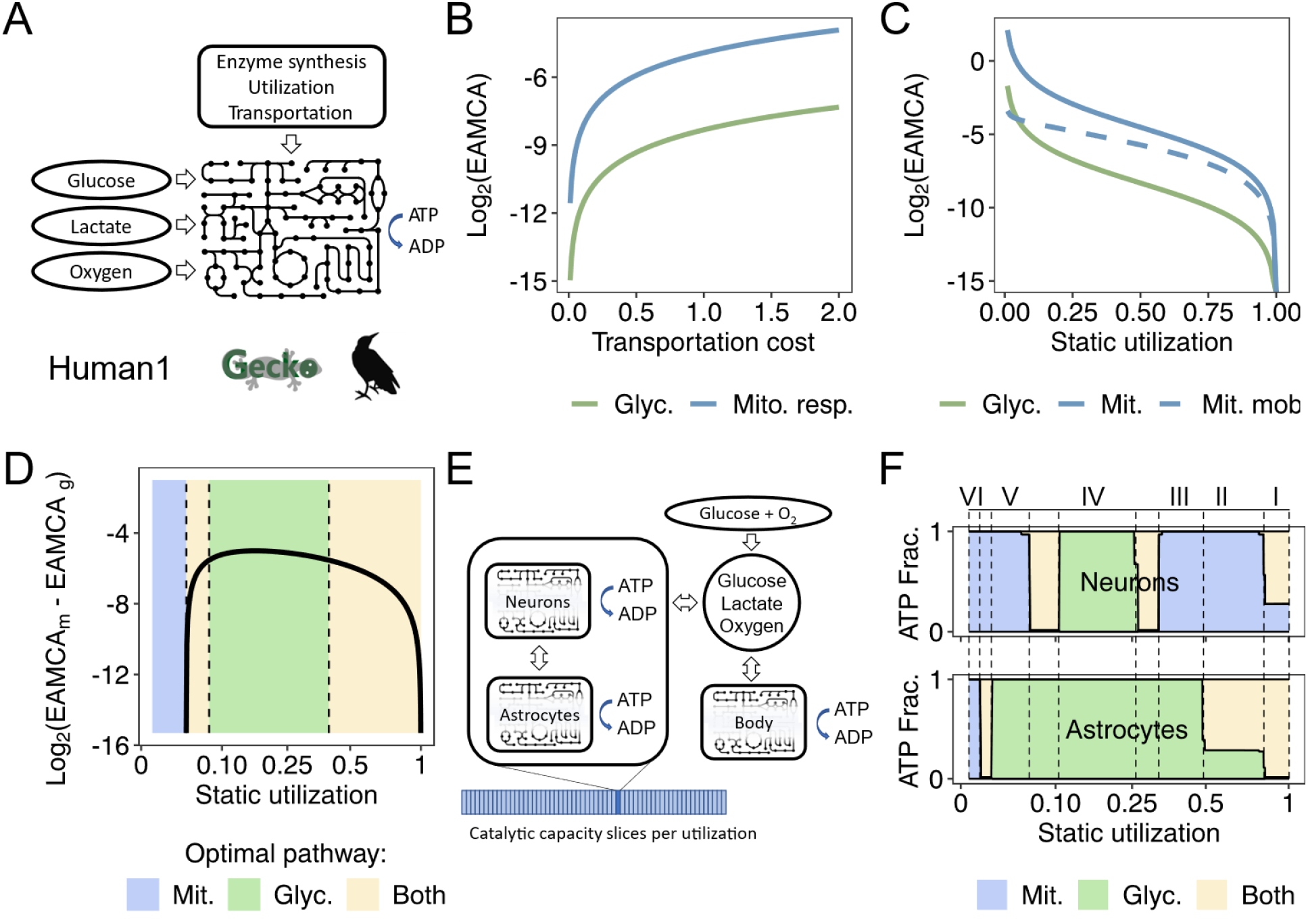
Investigation of brain energy metabolism using genome-scale metabolic models. A. Single neuron model. The full genome-scale model Human1 was extended with EAMCA for glycolysis and mitochondrial respiration. B. EAMCA per levels of transportation cost for each pathway. C. EAMCA per levels of utilization for each pathway. For the “Mit. Mob.” dashed line, the catalytic capacity of mitochondrial respiration is assumed to be used somewhere else 40% of the time it is not required at this specific location due to mitochondrial mobility. D. Reduction in EAMCA from using glycolysis instead of mitochondrial respiration at different levels of utilization, assuming mitochondrial mobility (as in Fig. 2C). The optimal pathway is chosen for glycolysis to cover the static utilization region where the reduction is highest, but the size of the glycolytic region is arbitrarily chosen to have the right border at 0.4. For simplicity, we have assigned both pathways to the block of utilization levels where glycolysis is beneficial and where we can in total have no lactate export. However, the optimal distribution between the pathways within this block is likely uneven, favoring glycolysis at high utilization levels. The x axis is transformed to Log_2_(U_s_ + 0.1). E. Combined model system simulating neurons, astrocytes, and the rest of the body. One astrocyte and neuron model exist per slice of static utilization, while the rest of the body is represented by a single model. Glucose and oxygen are freely distributed between the cell type models, while lactate transport is constrained. The different cell types are based on the neuron model but with different EAMCA parameters; the body model does not have any utilization or transportation costs while the utilization varies across utilization slices. F. Simulation results from using the combined model where astrocyte models are configured to have a lower utilization of mitochondria compared with neurons. The model predicts 6 utilization regions with different behaviors: I) Neurons help the body to clear lactate from the blood where EAMCA is low; II) ANLS and lactate uptake from blood in neurons (as in I); III) ANLS and export of lactate from astrocytes to the body. IV) Export of lactate from brain to body; V) ANLS and export of lactate from astrocytes to the body (same as III). VI) Uptake of lactate in both neurons and astrocytes. Regions of utilization where both pathways are used appear at switches between glycolysis and mitochondrial respiration – the use of both pathways in the same cell reduces the use of lactate conversion and transportation enzymes, and therefore reduces the enzyme maintenance costs given similar EAMCA for both pathways.

We used the model to estimate the EAMCA at different levels of utilization and transportation costs (Fig. 2B-C, Methods). Except for at very low utilization values, the EAMCA was higher for mitochondrial respiration compared with glycolysis due to the larger (~13.9 times) enzyme mass required for this pathway, despite simulated mitochondrial mobility. While the most optimal behavior would be to almost exclusively use glycolysis in the brain and consume the exported lactate in the rest of the body^17^, the brain would then produce more lactate than the body would be able to process. Glycolysis only yields 2 ATP per glucose and mitochondrial respiration generates roughly 29.5 ATP from the same amount of glucose (values derived from the model). We assume there is a limit to lactate production in the brain that is substantially lower than the theoretical maximum due to several reasons such as limiting the acidity in the blood and limited glucose supply. It is therefore important to use glycolysis in a way that optimizes the reduction in EAMCA. Assuming the same transportation costs for cytosolic and mitochondrial enzymes, glycolysis is most optimal to use at locations where transportation costs are high, such as at synapses far from the soma. With respect to utilization costs the optimization is more complex due to mitochondrial mobility, where the optimal allocation of glycolysis is at a low utilization range (not including the very lowest range) (Fig. 2D). These results fit well with observation 1 above: during high activity in a certain region of the brain, which engages catalytic capacity slices with low static utilization, the consumed oxygen vs. glucose ratio has been reported to decrease^1–4^, suggesting lactate export and thereby confirming our modeling results. Interestingly, one study showed that the ratio decreased the most directly after stimulation, and later became less attenuated^2^, suggesting that mitochondria move to synapses from other areas to help with ATP production and thereby increase the utilization of mitochondrial enzymes. In addition, different regions of the brain have been reported to exhibit different ratios of consumed oxygen vs. glucose^4^, which could potentially be an optimal behavior driven by different levels of static utilization, mitochondrial utilization, and enzyme transportation costs across such regions.

The ANLS makes lactate available to neurons, which can be useful due to the unsynchronized capacity of glycolysis and mitochondrial respiration (Fig. 1C). However, there might be a more important reason for its existence. Like neurons, astrocytes are spindle-shaped and are reported to stretch considerable lengths^18^; thus lactate production in astrocytes could be more optimal if the enzyme maintenance costs are different in such cells. Mitochondria are reported to be less mobile in astrocytes^19^, which could yield a lower utilization for mitochondria. Furthermore, while we have thus far modeled transportation costs equally for mitochondrial and cytosolic enzymes, the former could have lower transport costs since mitochondria are hypothesized to cycle to the soma for maintenance^20^. Both these effects were confirmed to induce ANLS using a metabolic model system of astrocytes and neurons (Fig. S2). This effect combined with varying EAMCA across the utilization range and a limited capability of the body to take up lactate results in a complex optimal enzyme allocation pattern. We generated a combined model system consisting of utilization slices containing neurons and astrocytes, which in turn were connected to the rest of the body via the blood, allowing for transport of lactate between cell types (Fig. 2E, Note S2). The combined model system configured with lower mitochondrial utilization in astrocytes and optimized to minimize the total enzyme maintenance cost yielded a complex optimal behavior across the utilization range, predicting both the ANLS theory and lactate export at high brain activity (Fig. 2F). A simulation with lower transportation costs for mitochondrial enzymes in neurons yielded similar results (Fig. S3).

Currently, little is known of the enzyme maintenance costs in the different cell types in the brain. From this work, we conclude that such information could help explain metabolic behaviors of the brain, opening a new avenue to explore in brain metabolism.

## Methods

### Model preparation and analysis

We used the full genome-scale metabolic model Human1^13^ v. 1.12 for model simulations. The model was enhanced by a modified version of GECKO Light^16,21^, which was used to assign enzyme usage ATP costs to reactions based on *k_cat_* values and molecular weights. The costs were divided into 3 categories that could be penalized differently: 1) MT genes (i.e., enzyme subunits generated from the mitochondrial genome), 2) other mitochondrial enzymes (i.e., enzymes or enzyme subunits that reside in the mitochondria but are generated in the cytoplasm of the soma and thus need to be transported to the mitochondria at the synapses), and 3) other enzymes (including for example cytosolic enzymes). All model simulations were performed using flux balance analysis as implemented in the RAVEN Toolbox^22^.

### Modeling ATP maintenance costs

We generated a generic cell type model to which ATP costs from enzyme usage could be added. This was done by adding pseudo-metabolites. Thus, each enzyme catalyzed reaction was set to consume different quantities of three different pseudo-metabolites representing enzyme-usage of three categories: MT-genes, other mitochondrial enzymes, and other enzymes. For each pseudo-metabolite, a maintenance reaction was added, which produces the pseudo-metabolite and hydrolyses ATP to simulate a maintenance cost. The maintenance cost can be configured differently for the different enzyme categories by varying the stoichiometric coefficient for the pseudo-metabolite. Each maintenance reaction has the following form:

H2O + ATP => ADP + Pi + H^+^ + S_x_ x_prot_pool

where x_prot_pool is the pseudo-metabolite of the enzyme-usage category *x* and *S_x_* is the stoichiometric coefficient by which the metabolite for category *x* (*mt, me*, or *oe*) is produced. *S_x_* can be derived from the protein maintenance cost *C_x_* as

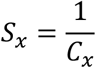

The maintenance cost for each category can in turn be derived from the utilization of glycolysis (*U_g_*) and mitochondrial respiration (*U_m_*, here representing both the MT genes and other mitochondrial enzymes), the transportation cost, and a basic maintenance cost *B*, which represents the ATP cost of maintaining one unit of protein usage (set to 1 mmol ATP gDW^-1^h^-1^ in all our simulations unless otherwise specified – the value has no impact on the qualitative results). The absolute transportation cost *T_a_* is assumed to be proportional to *B* with the proportionality constant *T_x_*, which we for convenience call the transportation cost:

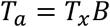

In our simulations, *T_x_* ranges between 0.01 and 2, with a default value of 0.1 unless otherwise specified, and is assumed to be the same for mitochondrial respiration and glycolysis (we assume both categories are transported using fast transport^9^, where glycolytic enzymes are mainly transported in vesicles^23^). However, protein subunits generated from MT genes are assumed to be produced locally in mitochondria and therefore have a transportation cost of 0.

The total protein maintenance cost of enzyme category *x* is defined as

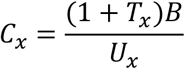

The division by the utilization is motivated as follows: We assume that all enzymes need to be maintained regardless of their usage and ignore any effect on enzyme half-life from enzyme usage. The maintenance cost per enzyme usage will then be proportional to *U_x_*^-1^ since the maintenance cost for the time periods the enzyme is not used is accounted for during the time periods it is in use.

To estimate the difference in protein use per pathway and ATP produced we used the model without enzyme maintenance costs. For glycolysis, the full Human1 model was given unlimited access to H_2_O, phosphate, and H^+^, and in addition 1 mmol^*^gDW^-1^h^-1^ of glucose, but no oxygen, while we supplied unlimited oxygen and lactate instead of glucose for the mitochondrial respiration. In both cases, we ran FBA optimized for maximum flux through an ATP hydrolysis reaction. The ratio *R* between the enzyme mass usage per ATP was calculated as

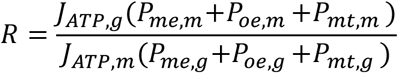

where *J_ATP,g_* and *J_ATP,m_* are the ATP production fluxes for glycolysis (*g*) and mitochondrial respiration (*m*) (equal to the flux through the maintenance reaction in the simulations), and *P_x,g_* and *P_x,m_* are the corresponding protein usages in the two pathways for the three different enzyme categories. *R* was estimated to 13.91 from the model simulations.

To estimate the EAMCA for Fig. 2B-D we ran similar simulations but with extra maintenance costs. The EAMCA was then calculated as the difference between a run with no transportation and utilization costs and a run with such extra maintenance costs included. Since the modeling setup does not allow for an EAMCA > 1, we reduced the value of B to 0.1 mmol ATP gDW^-1^h^-1^ in the simulations (Mitochondrial respiration can get EAMCA > 1 for very low utilizations, which happens in Fig. 2C), and compensated this trick by multiplying the resulting EAMCA by 10, which yields the same results as what would be expected from setting B to 1 mmol ATP gDW^-1^h^-1^ in the simulations. The transportation cost was set to 0 in Fig. 2C and 0.1 in Fig. 2D, and the utilization was set to 1 in Fig. 2B.

### Modeling maintenance reduction from mitochondrial mobility

The catalytic capacity utilization for the mitochondrial respiration pathway when taking mitochondrial mobility into account, *U_m,m_*, was assumed to be

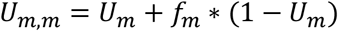

where *f_m_* is a “mobility factor” set to 0.4 in our simulations. (1 - *U_m_*) is the time fraction the marginal catalytic capacity investigated is not used at the physical position of interest and *f_m_* is the assumed fraction of this time that the marginal enzymatic capacity can be used somewhere else. We have not included any ATP costs for mitochondrial movement in this model since this is difficult to estimate.

### Combined models

The combined model used in Fig. S2 is described in Fig. S2A. The combined model used in figure 2E-F is described in detail in Note S1.

### Acquisition of enzyme half-life data

Enzyme half-life data for HELA cells were obtained from the literature^24^. All proteins that had a matching gene association for any reaction used in each pathway (glycolysis or mitochondrial respiration) were deemed to be associated with the pathway and included in Fig. S1.

### Software

The data was analyzed using MATLAB R2019b and R version 3.6.1. To ensure the quality of our analyses, we verified and validated the code using a combination of test cases, reasoning around expected outcome of a function, and code review. The details of this activity are available in the verification matrix available with the code.

## Declarations

### Availability of data and materials

The model Human1 is available in GitHub (https://github.com/SysBioChalmers/Human-GEM).

The processed data and source code are available in Zenodo: https://doi.org/10.5281/zenodo.7320333. The source code is also available in GitHub: (https://github.com/SysBioChalmers/BrainMetabolismModeling).

### Authors’ contribution

J.G came up with the original idea. J.G, J.R, and J.N. planned the project. J.G. wrote all software and made all analyses and figures. J.G. wrote the draft manuscript. All authors reviewed and edited the manuscript. J.R. and J.N. supervised the project. J.N. acquired funding for the project.

## Acknowledgements

None.

