## Supplementary information for "Brain energy metabolism is optimized to minimize the cost of enzyme synthesis and transport"

* Corresponding author

### Supplementary Figures

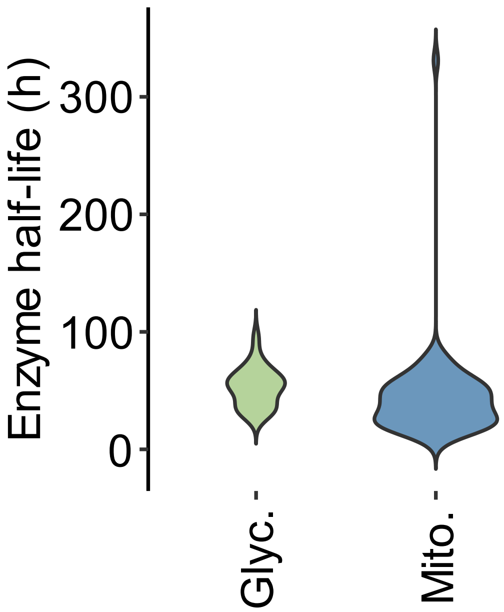


***Fig S1: Difference in enzyme half-lives for enzymes participating in glycolysis and mitochondrial respiration.***

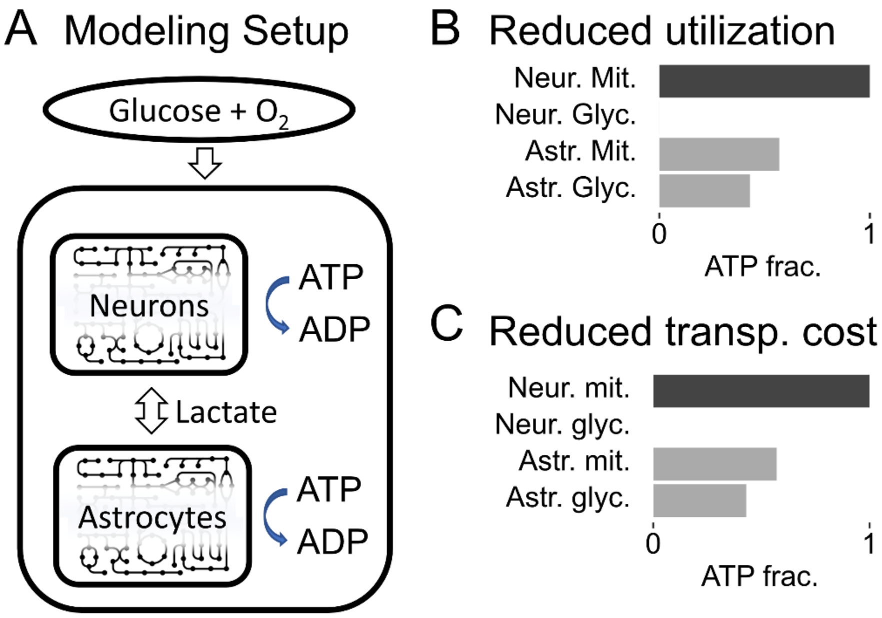


***Fig S2: Modeling of neurons and astrocytes.*** *A. Modeling setup. Two models representing neurons and astrocytes were given glucose and oxygen as input. In addition, lactate was allowed to be transported freely between the cell types. The model system was optimized to maximize flux through an ATP hydrolysis reaction, where 15% of the ATP was set to be hydrolyzed in the astrocytes and the rest in the neurons. The models support the addition of maintenance costs from enzyme utilization and transportation. B. Simulation where the mitochondrial utilization was reduced in astrocytes due to less mitochondrial mobility in such cells. The mitochondrial mobility factor (see Methods in the main text) was set to 0.4 for neurons and 0.2 in astrocytes. For both cell types, the static utilization was set to 0.5 and all transportation costs were set to 0.1. The x axis represents the fraction of ATP generated per pathway within each cell type. C. Simulation where the mitochondrial transportation was lower in neurons due to increased mitochondrial mobility in such cells (see main text). Specifically, the transportation cost for mitochondrial enzymes in neurons was set to 0.1, while all other transportation costs were set to 0.2. for both cell types. For both cell types, the mitochondrial mobility factor was set to 0.4 and the static utilization was set to 0.5. Strikingly, the results of the simulations in B and C are very similar - all glycolysis take place in the astrocytes in both cases.*

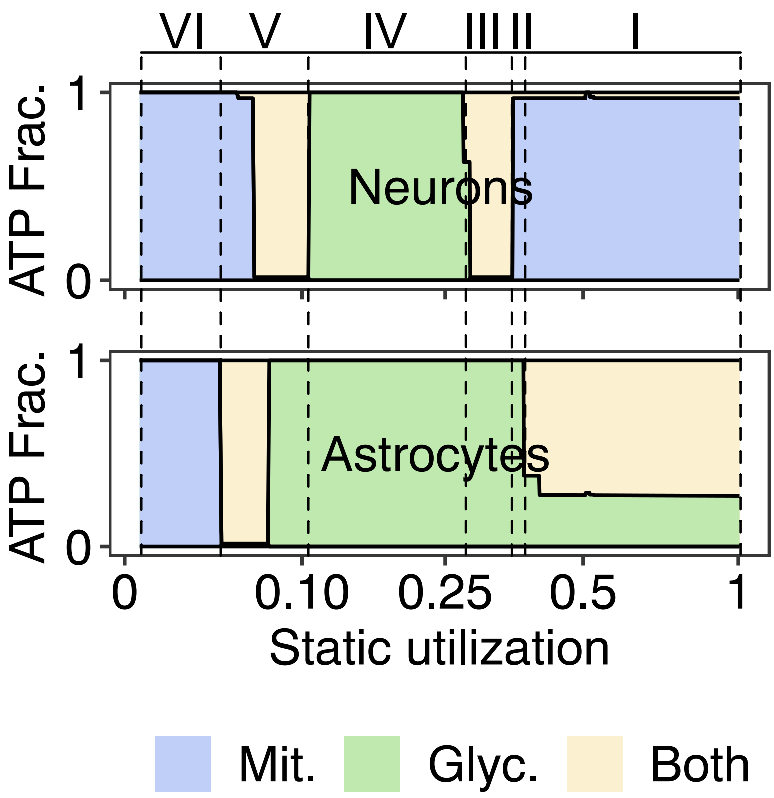


***Fig S3: Simulation results from using the combined model system where neuron models are configured to have a lower transportation cost for mitochondrial enzymes compared to astrocytes.*** *The model system (as described in Fig. 2E in the main text) predicts 5 utilization regions with different behaviors: I) ANLS and lactate uptake by neurons from the blood; II) ANLS and export of lactate from astrocytes to the body. III) Gradual change from mitochondrial respiration to glycolysis in neurons. IV) Export of lactate from brain to body; V) Gradual change towards import of lactate in both cell types; the change happens first in neurons and later in astrocytes. VI) Import of lactate to both cell types.*

### Supplementary Notes

#### Note S1 – The car transport analogy

To further explain the concept of catalytic capacity utilization, we here provide an example of transport using cars as an analogy. In this analogy we envision a transportation company that needs cars to perform their work. However, the number of cars they need varies from day to day (Fig. I A). There are two ways to supply those cars: to own them at a cost of $30 per day, or to rent them at a cost of $90 per day. However, while it is necessary to pay for owned cars every day (regardless of their use), renting cars allows more flexibility and does not require payment on days when such cars are not needed. In this example, we introduce car slots (numbered 1-8), where each slot represents an available position in the fleet, which can be occupied either by an owned or rented car (Fig. I B). The car slots are always used from left to right.

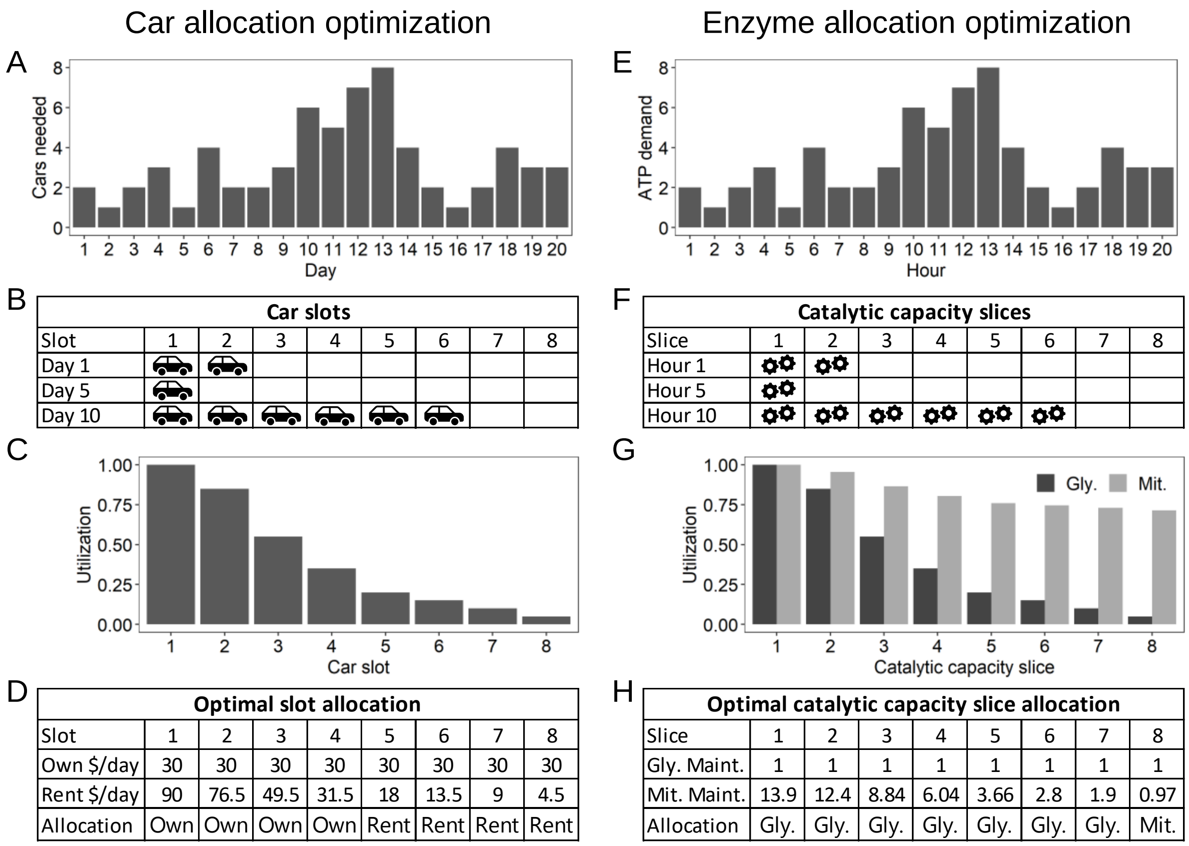


| Optimal slot allocation | | | | | | | | |
|---|---|---|---|---|---|---|---|---|
| Slot | 1 | 2 | 3 | 4 | 5 | 6 | 7 | 8 |
| Own $/day | 30 | 30 | 30 | 30 | 30 | 30 | 30 | 30 |
| Rent $/day | 90 | 76.5 | 49.5 | 31.5 | 18 | 13.5 | 9 | 4.5 |
| Allocation | Own | Own | Own | Own | Rent | Rent | Rent | Rent |

| Optimal catalytic capacity slice allocation | | | | | | | | |
|---|---|---|---|---|---|---|---|---|
| Slice | 1 | 2 | 3 | 4 | 5 | 6 | 7 | 8 |
| Gly. Maint. | 1 | 1 | 1 | 1 | 1 | 1 | 1 | 1 |
| Mit. Maint. | 13.9 | 12.4 | 8.84 | 6.04 | 3.66 | 2.8 | 1.9 | 0.97 |
| Allocation | Gly. | Gly. | Gly. | Gly. | Gly. | Gly. | Gly. | Mit. |

***Fig. I The car transport analogy.*** *A. Cars needed per day (fictive data). B. Use of car slots per day for a few example days. C. Utilization of each car slot. D. Average daily costs for the two car allocation strategies given the utilization, followed by the optimal slot allocation. The cost is calculated as the average cost per day across all 20 days. E. ATP demand per hour (mmol ATP gDW$^{-1}$h$^{-1}$, fictive data). F. Use of catalytic capacity slices for a few example hours. G. Utilization of each catalytic capacity slice for the two pathways, where we assume that mitochondria can be used somewhere else 70% of the time they are not needed at this specific synapse. H. Relative ATP maintenance costs for each slice for the two pathways, taking utilization into account, followed by the optimal pathway allocation for the slices. The cost is relative to the cost of producing 1 mmol ATP gDW$^{-1}$h$^{-1}$ using glycolysis.*

We then calculate how much each car slot is used, which we define as the utilization (*U*) of each slot, where the utilization is the fraction of the days a slot is used. For example, the first car slot will be used every day and will hence have a utilization of 20/20 = 1, while the 8$^{th}$ slot will only be used one day out of twenty, resulting in a utilization of 1/20 = 0.05. To optimize the cost for the options of owning vs renting cars for the different slots, we can calculate the total cost per day for each slot across the 20 days. For owned cars it is a constant cost per slot per day ($30), while the average cost per slot per day for renting a car, $C_{r,\,avg}$, can be calculated as

$$C_{r,avg} = UC_d$$

where $C_d$ is the daily cost for renting a car ($90). Considering the utilization of the car slots, the optimal behavior is thus to own 4 cars and rent the remaining cars when needed (Fig. I D).

We next turn our attention to catalytic capacity utilization. We here have a varying ATP demand per hour at a fictive synapse and use the same data distribution as for the car allocation case (Fig. I E). While we in the real case have an (almost) infinite number of catalytic capacity slices, we here for simplicity divide the total capacity into 8 slices each with the capacity to produce 1 mmol ATP $gDW^{-1}h^{-1}$. The slices are used in a similar manner as the slots in the car case (Fig. I F). The optimization problem is similar, where we need to decide in advance to which pathway each slice is allocated. The utilization works a bit differently – here mitochondria can be shared with other physical locations in the cell, such as other synapses. The utilization for mitochondrial respiration is thus higher than that of glycolysis, and we in this case assume a mobility factor of 0.7 (see Methods), meaning that the mitochondria can be used somewhere else 70% of the time they are not used at this synapse (Fig. I G).

To optimize the pathway allocation to the different slices, we next calculate the total relative maintenance cost of each slice for the two pathways. The cost of glycolysis is constant (similar to the case where the company owns a car) while the cost of mitochondrial respiration varies with the utilization, since we assume that the maintenance costs can be shared with other physical locations in the cell. The fraction *f* of the maintenance costs that need to be paid by this synapse can then be calculated as

$$f = \frac{U_m}{(U_m + 0.7(1 - U_m))}$$

where $U_m$ is the utilization for mitochondria. The optimal allocation of pathway to the slices is in this case to assign glycolysis to the first 7 slices and mitochondrial respiration to slice 8 (Fig. I H).

### Note S2 – Detailed description of the full body model

A full body model system was used to simulate how the optimal metabolism varies across the utilization range in a brain model consisting of astrocytes, neurons, and the rest of the body. Of the total ATP hydrolyzed in the body, the astrocytes were assumed to hydrolyze 3%, the neurons 17%, and the rest of the body 80%. While the ATP need of neurons and astrocytes is spatially distributed both within the cells but also across cells and brain regions, we have as a simplification assumed that the transportation cost (set to 0.1 for astrocytes and neurons) and distribution of utilization is equal across all such spatial points, allowing for modeling all such points together. The neurons and astrocytes were therefore modeled in pairs of one cell type each, where each such pair represented a catalytic capacity slice at a certain static utilization (Fig. 2E in the main text). We modeled in total 100 such slices, creating a combined model with 201 entity models with static utilization slices ranging from 0.01 to 1. The rest of the body model has no extra maintenance costs from transportation or utilization.

The models were configured according to the following rules:

1. For simplicity, we assumed that each utilization slice contained an equal ATP production capacity, although it is not equally used in all slices.
2. All cell models had unlimited access to glucose and oxygen, although the total glucose uptake rate was restricted to 1 mmol $gDW^{-1}h^{-1}$.
3. A combined ATP hydrolysis reaction was generated, which was used as objective function in the simulation. 80% of the ATP was taken from the rest of the body, 17% from neurons, and 3% from astrocytes. For the neurons and astrocytes, the ATP was taken from each slice such that the ATP taken from each slice was proportional to their static utilization since each slice only produces ATP when in active use.
4. Lactate could be freely exchanged within a utilization slice, which enables lactate from astrocytes to flow to neurons.
5. Lactate transport into the slices and the rest of the body was restricted to 50% of the glucose transport required when all ATP is generated from glucose, in both the slices and the rest of the body. In practice, this constrains the ATP production from lactate from blood in slices and the rest of the body from exceeding ~25% of the total ATP production, although export between astrocytes and neurons is unrestricted. The motivation for this constraint is that the concentration of lactate is substantially lower than that of glucose in blood.
6. Lactate can be freely transported to a slice from other slices with a higher utilization. This is motivated by that when the enzymatic capacity of a slice is in use, all slices with higher utilization are also in use. In practice, these fluxes are small, and there is therefore no need to constrain them. These fluxes help feed the neurons in region VI in Fig. 2F in the main text.
7. For neurons and astrocytes we assumed that the mitochondria are shared across physical locations within cells, making it possible to share maintenance costs for mitochondrial enzymes, reducing the time a slice was not used (caused by static utilization) by 40%. This was simulated by setting the mobility factor to 0.4 (see Methods).

In Fig. 2F, the mobility factor for astrocytes was set to 0.2, which increases the EAMCA caused by utilization for mitochondrial respiration in astrocytes.

In Fig. S3, we set the transportation costs of mitochondrial enzymes to 0.1 in neurons and other transport costs to 0.2. The mobility of mitochondria was in this case assumed to be able to reduce the transportation cost of enzymes to mitochondria, since it may be cheaper to move whole mitochondria closer to the soma for enzyme replacement than moving many vesicles with enzymes to and from the synapses.

Although all spatial positions are modeled together, we don't expect that the ATP need is synchronized across such points – when there is a peak ATP demand at one point, another may be at rest. It is therefore reasonable to assume that points with a low activity (with only slices with high utilization active) may use lactate produced at points with high activity, where the lactate is transported via the blood. In practice, this is represented in the model by lactate uptake from the blood at high utilization slices (which are used at rest) and lactate export from low utilization slices (which are used at peak hours).

To make it practically possible to combine 201 copies of the Human1 model, we first minimized the model, only including reactions that carried flux in any of 3 simulations all optimized for ATP hydrolysis: 1) The model was fed on glucose; 2) The model was fed on glucose and oxygen; and 3) The model was fed on lactate and oxygen. The resulting reduced model had 55 reactions in total.
